# Morphological deconvolution of beta-lactam polyspecificity in *E. coli*

**DOI:** 10.1101/545335

**Authors:** William J. Godinez, Helen Chan, Imtiaz Hossain, Cindy Li, Srijan Ranjitkar, Dita Rasper, Robert L. Simmons, Xian Zhang, Brian Y. Feng

## Abstract

Beta-lactam antibiotics comprise one of the earliest known classes of antibiotic therapies. These molsecules covalently inhibit enzymes from the family of penicillin-binding proteins, which are essential to the construction of the bacterial cell wall. As a result, beta-lactams have long been known to cause striking changes to cellular morphology. The exact nature of the changes tend to vary by the precise PBPs engaged in the cell since beta-lactams exhibit a range of PBP enzyme specificity. The traditional method for exploring beta-lactam polyspecificity is a gel-based binding assay which is low-throughput and typically run *ex situ* in cell extracts. Here, we describe a medium-throughput, image-based assay combined with machine learning methods to automatically profile the activity of beta-lactams in *E. coli* cells. By testing for morphological change across a panel of strains with perturbations to individual PBP enzymes, our approach automatically and quantifiably relates different beta-lactam antibiotics according to their preferences for individual PBPs in cells. We show the potential of our approach for guiding the design of novel inhibitors towards different PBP-binding profiles by recapitulating the activity of two recently-reported PBP inhibitors.

## Introduction

Beta-lactam antibiotics are a crucial part of the therapeutic arsenal. These molecules inhibit bacterial cell wall biosynthesis, and are efficacious against a broad spectrum of bacterial pathogens. Over the last few decades, researchers have systematically modified the beta-lactam scaffold, seeking to evade recognition by beta-lactamase enzymes which bacteria use to inactivate these drugs. Spread of the genes encoding beta-lactamases limits the clinical utility of beta-lactam antibiotics. The development of this family of molecules and the resulting evolutionary selection of beta-lactamases exemplifies the chemical arms race between bacteria and mankind.^1^

Beta-lactams have also served as critical tools in the study of bacterial cell wall biosynthesis. These molecules stably acylate a family of enzymes which were named penicillin-binding proteins (PBPs). These enzymes coordinate cell-wall biosynthesis in an intricate process involving multiple multi-protein complexes (recently reviewed by Dorr and colleagues).^2^ Numerous PBP enzymes have been identified in bacteria, and they can be generally classified according to their enzymatic function. In *E. coli*, class A PBP’s, such as PBP1a and PBP1b, are bifunctional enzymes that catalyze both transglycosylation (polymerization) of the peptidoglycan and transpeptidation (cross-linking) of the glycan strands. Class B enzymes such as PBP2 and PBP3 are monofunctional transpeptidases. Class C enzymes, which include PBP4, PBP4b, PBP5, PBP6, PBP6b, and AmpH carry out various peptidase reactions that facilitate maturation, remodeling, and metabolism of peptidoglycan.^3,4^ These activities are critical in cell elongation and division.

Beta-lactam molecules range in specificity, and our understanding of this polyspecificity primarily owes to gel-based competition-binding assays using labeled beta-lactam analogues.^5–10^ These assays are powerful because they provide simultaneous measurements on all penicillin-binding proteins in the extract. However, they are low-throughput and sensitive to many experimental variables such as extraction conditions, binding kinetics and dose, and as a result published values for binding affinities vary. Binding assays are also blind to cellular variables such as permeability and efflux, though some recent work has suggested the gel-based competition method can be applied to intact cells.^6^

Inhibition of PBP enzymes has also been shown to induce a diverse array of cellular morphologies, reflecting the roles they play in coordinating division and elongation.^11-13^ Inhibition of PBP2 and PBP3, which are essential for *E. coli* growth, result in filamentous and ovoid morphologies, respectively.^14-17^ PBP1a and PBP1b, which are not individually essential, form a synthetic lethal pair; loss of both induces cell lysis.^18^ Class C PBPs, though also not essential, have been linked to more subtle morphological effects.^19-22^ Recent work by Kishony and colleagues has also established the time-dependence of beta-lactam-induced morphological change, illustrating that different beta-lactams can cause chronologically distinct morphological changes.^23^ And though technologies such as flow cytometry and electron microscopy have been successfully used to study morphology^24^ the most common approach relies on light microscopy using high-magnification oil objectives and samples immobilized on a substrate such as agarose.^25^ Consequently, these methods have limited throughput.

To improve our ability to study beta-lactam activity in cells, we sought to develop a method that was both quantitative and amenable to medium- or high-throughput applications. Recent advances in automated microscopy have led to the field of high-content imaging (HCI),^26,27^ whereby high-throughput image acquisition is paired with automated image analysis methods to support cell-based screening. Though this approach is widely used in mammalian cellular assays,^28,29^ it has recently been explored for microbiological applications.^30-38^ However, difficulties immobilizing bacterial samples and image analysis challenges, such as clearly segmenting bacterial cell objects, have limited this approach.

In this work, we present an approach that combines medium-throughput imaging of *E. coli* cells at high-magnification with machine learning-based automated image analysis. We use this approach to quantitatively characterize the PBP-binding preferences of beta-lactam antibiotics based on the morphological variations they induce. Using methods based on deep neural networks^39,40^ and Hidden Markov models,^41,42^ we classify cellular morphologies and capture the progression of morphology over dose into compound-specific profiles. By comparing these profiles of morphological change across a panel of cell lines with single perturbations to PBP enzymes, we demonstrate how morphological profiles of beta-lactam antibiotics reflect their affinities for individual PBPs in cells. We show that this analysis compares favorably to both internal and published gel-based competition data and provides additional insights into the physiological effects of PBP inhibition. By applying the approach to historical and recent PBP inhibitors, we show the potential usefulness of the approach for guiding optimization of novel beta-lactams or PBP inhibitors.

## Results

### Building morphology profiles of reference b-lactams from bacterial HCS images via deep neural networks

PBP enzymes build PG in a highly interdependent fashion, and the process—which includes synthesis, cross-linking, cleavage and recycling roles—involves enzymes which may be at least partially redundant. Young and colleagues explored this concept explicitly by making every combination of non-essential PBPs. Strikingly, cells lacking PBPs 4, 5, 6, 7 and 8 are viable.^43^ To emphasize the individual roles of PBP enzymes, we assembled a panel of *E. coli* strains with perturbations to PBP function, hypothesizing that surveying such a panel would facilitate discrimination between promiscuous and specific beta-lactams. This panel was comprised of deletion strains for PBP1a, PBP1b, PBP4 and PBP5. Because PBP2 is essential, we studied the importance of this enzyme by including a strain of *E. coli* expressing a PBP2 mutant (PBP2R)^44,45^ less susceptible to some PBP2 inhibitors (**Supplementary Table 1**). *E. coli-ΔPBP5* was included because of its reported role in contributing to cellular morphology.^21,46-48^ *E. coli-ΔPBP4* was included as it has not been shown to have a strong role in defining cell morphology.^20,22^ To understand the range of beta-lactam PBP preferences, we assembled a collection of 12 beta-lactams spanning the major chemical classes (**Table 1**). We also included D-cycloserine, a dual inhibitor of D-ala-D-ala ligase and alanine racemase, which inhibits cell-wall biosynthesis upstream of the PBP enzymes.

**Table 1.** Compounds used in this study. *Primary target denoted by internal gel-based binding assays. Citations indicate primary target in reference literature. **DADAL/AR = D-ala D-ala ligase / Alanine racemase.

| Compound | Structure | Chemical Class | Primary Target* | Compound | Structure | Chemical Class | Primary Target* |
| --- | --- | --- | --- | --- | --- | --- | --- |
| ampicillin |  | penicillin | PBP3 | D-cycloserine |  | Serine analogue | DADAL/AR** |
| mecillinam |  | penicillin | PBP2 | faropenem |  | penem | PBP2 |
| piperacillin |  | penicillin | PBP3<sup>6</sup> | doripenem |  | carbapenem | PBP2<sup>6,51</sup> |
| cephalexin |  | 1<sup>st</sup> gen. cephalosporin | PBP3<sup>6</sup> | meropenem |  | carbapenem | PBP2 |
| cefoxitin |  | 2<sup>nd</sup> gen. cephalosporin | PBP 1A/1B<sup>6,52</sup> | aztreonam |  | monobactam | PBP3 |
| cefsulodin |  | 3<sup>rd</sup> gen. cephalosporin | PBP 1A/1B<sup>6,52</sup> | carumonam |  | monobactam | PBP3<sup>53</sup> |
| moxalactam |  | 3<sup>rd</sup> gen. cephalosporin | PBP3<sup>54</sup> | BAL30072 |  | sulfactam | PBP3<sup>55</sup> |
|  |  |  |  | NXL105 |  | Diazabicyclooctane | PBP2 |

Each cell line in the panel was treated with the collection of beta-lactams at 8 concentrations, standardized relative to the MIC of each compound in each cell line (**Supplementary Table 1**). Samples were prepared for imaging by treating exponential-phase cells with compounds for two hours, followed by simultaneous fixation and staining with fluorescent markers for cell membrane (FM4-64FX, Invitrogen), nucleic acids (Syto9, Invitrogen), and a DNA-specific stain (Hoechst 34580, Life Technologies). Cells were imaged in 96-well plates at 100X magnification.

Treatment led to four cellular morphologies precedented in the PBP literature: ovoid, filament, spindle, and lysed.^49,50^ In addition, we considered the untreated morphology and an enlarged morphology that commonly appeared in treatments with low compounds concentrations (**Fig. 1**). We reasoned that this phenotype may be a transitional morphology between untreated and the more canonical morphologies caused by PBP inhibitor treatment.

**Figure 1.**
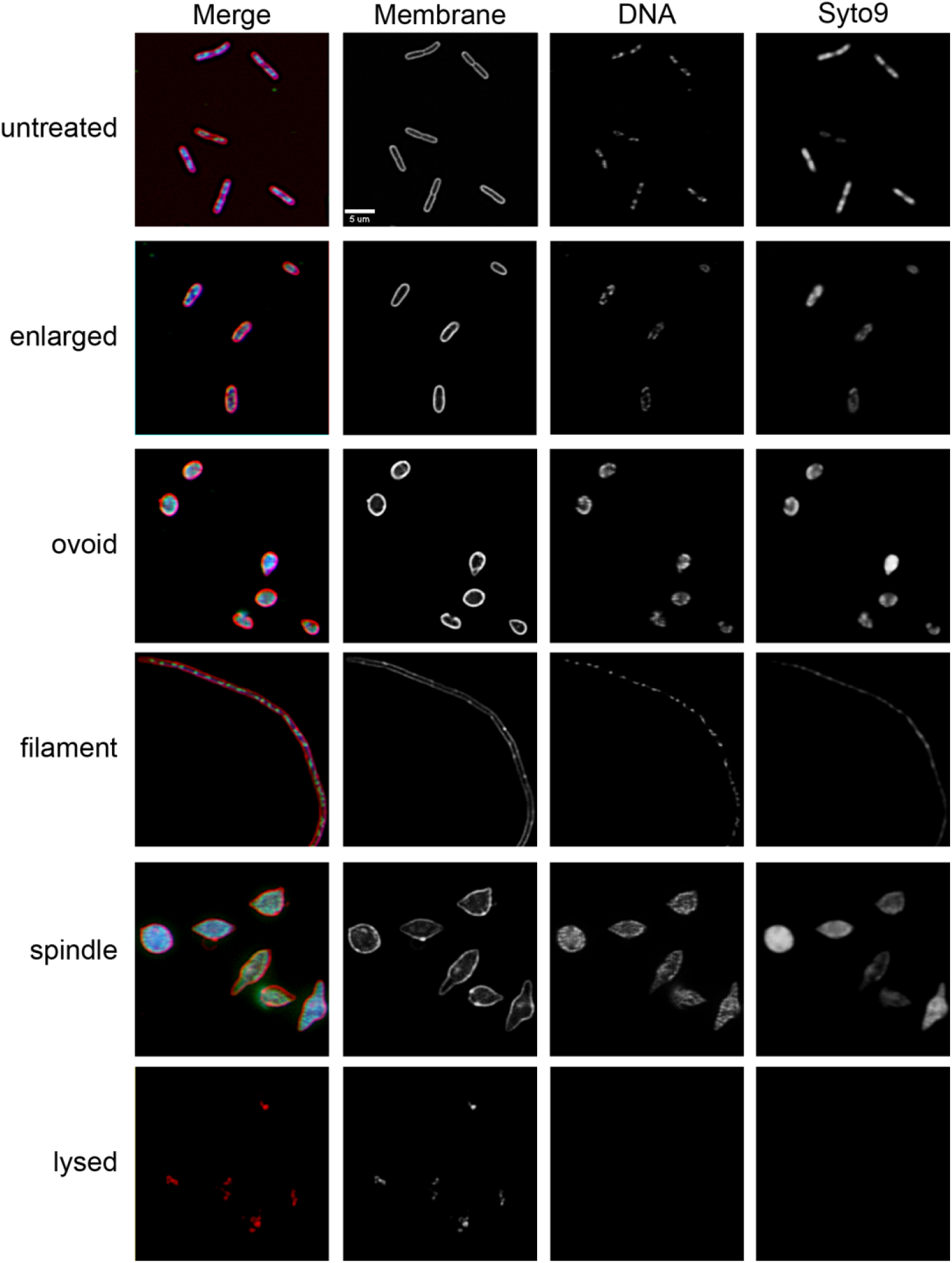
Representative images of each morphological class. Membranes stained with FM4-64FX. DNA was stained with Hoechst 34580. Nucleic acids were stained with Syto9. Images were acquired at 100X magnification.

To automatically classify images displaying multiple bacteria into these six categories, we used a supervised machine learning^39^ approach based on a multi-scale convolutional neural network (M-CNN) architecture^40^ that takes as input a full-resolution three-channel fluorescence microscopy image and yields as output a probability score for each of the six morphologies. We took the morphology category with the largest score as the morphology prediction for the input image (see workflow in **Fig. 2a**). All model parameters were optimized automatically using 129 images (and variations thereof obtained through data augmentation strategies) annotated manually with one of the six available morphologies (see **Materials and Methods**).

**Figure 2.**
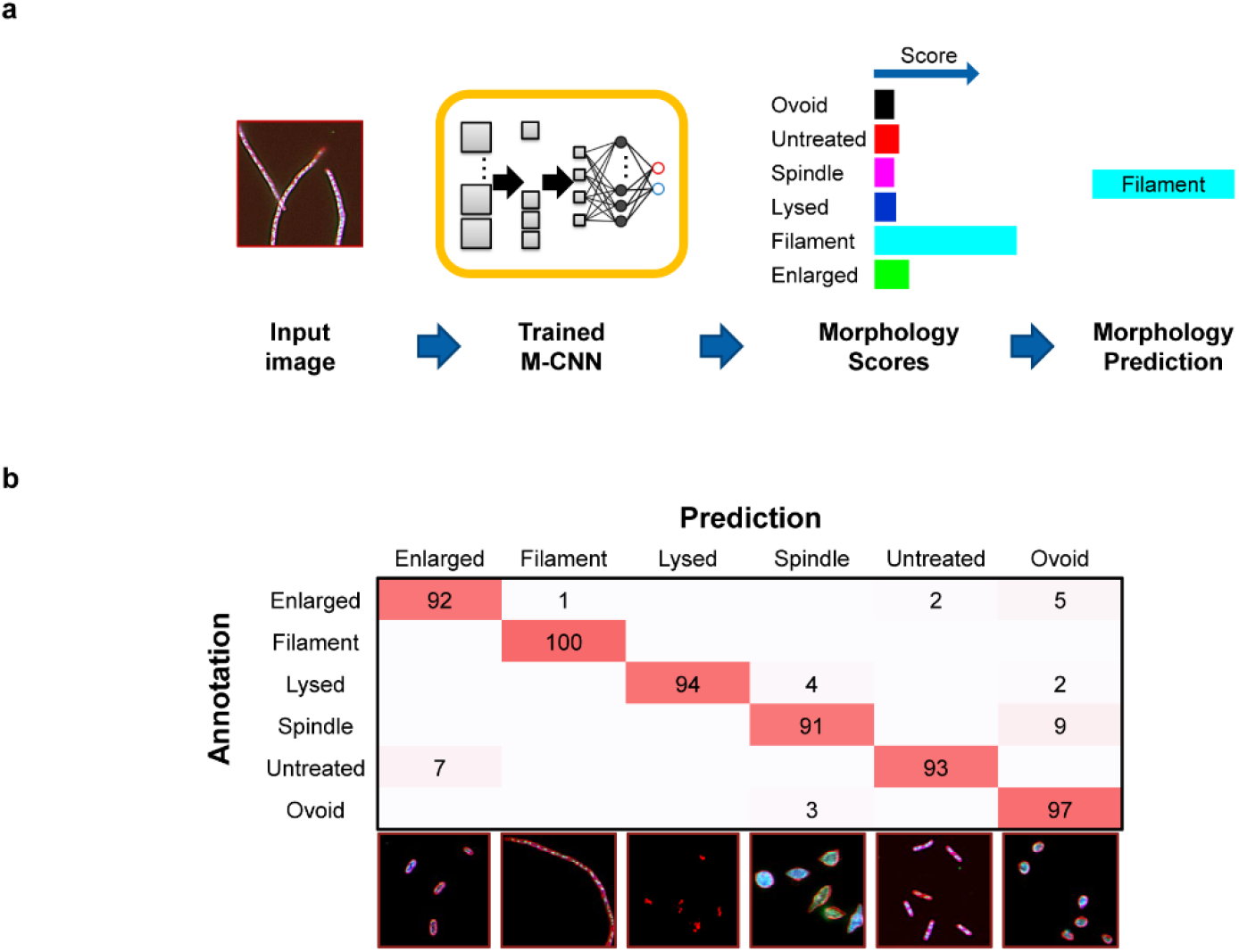
From pixels to morphologies with deep learning. (**a**) Taking as input a multi-channel multi-cellular image, the trained multi-scale convolutional neural network (M-CNN) model yields a probability score for each of the six morphologies included in the model. The morphology with the largest score is taken as the morphology prediction for the entire image. (**b**) Classification accuracy given in percentage terms [%] for the M-CNN model evaluated through cross-validation. Each row of the displayed confusion matrix shows the true morphology annotation while the columns show the predictions from the M-CNN architecture. Entry values are normalized so that the sum of each row over the columns add up to a value of 100%. Entries are shaded in red and the shading intensity correlates with the relative magnitude of the values. Entries without numbers indicate values of 0. Example three-channel images (cell membrane, red; DNA, green; nucleic acid, blue) from each class are shown below the table.

To validate the classification accuracy of the method, we applied a repeated random holdout cross-validation approach where we randomly selected 90% of the manually annotated images of each morphology category to optimize the model parameters (training set). Once optimized, we applied the network model to the remaining 10% of the images of each morphology category (test set), and recorded the network’s morphology predictions for those images. We repeated this process 50 times, and compared the network’s predictions with the true morphology labels. The confusion matrix in **Fig. 2b** shows the results of this comparison. The diagonal entries of the matrix show the fraction of images correctly classified for each morphology category. The off-diagonal entries show the fraction of images that were misclassified. A perfect model would yield 100% on all diagonal entries and 0% on all off-diagonal entries. Our model yields values above 90% on all diagonal entries. The highest (100%) accuracy is obtained on images from the filament category, while the lowest accuracy (91%) is obtained on images from the spindle category, which are occasionally confused with the ovoid category. Over all morphologies, the classification accuracy is 95 ± 2.57% (mean ± 95% confidence interval).

Once we had ascertained the validity of the M-CNN approach for identifying PBP morphologies, we applied an M-CNN model optimized on all available annotated images to 14822 full-resolution images displaying the morphological outcomes induced by compounds at eight multiples of the minimal inhibitory concentration (MIC) across all six cell lines (predictions for all images are provided in **Supplementary Data)**. The morphology predictions of images belonging to a certain experimental combination of compound, fold MIC, and cell line were collapsed onto a single morphology prediction (see **Materials and Methods**). For each compound, we arranged the morphology predictions onto a color-coded 2D matrix over increasing MIC folds (x-axis) and cell lines (y-axis). Color-coded morphology profiles of reference beta-lactams are shown in Fig. 3. These visualizations, which we colloquially refer to as ‘Tetris’ plots, serve as simple approximations of the model’s output.

**Figure 3.**
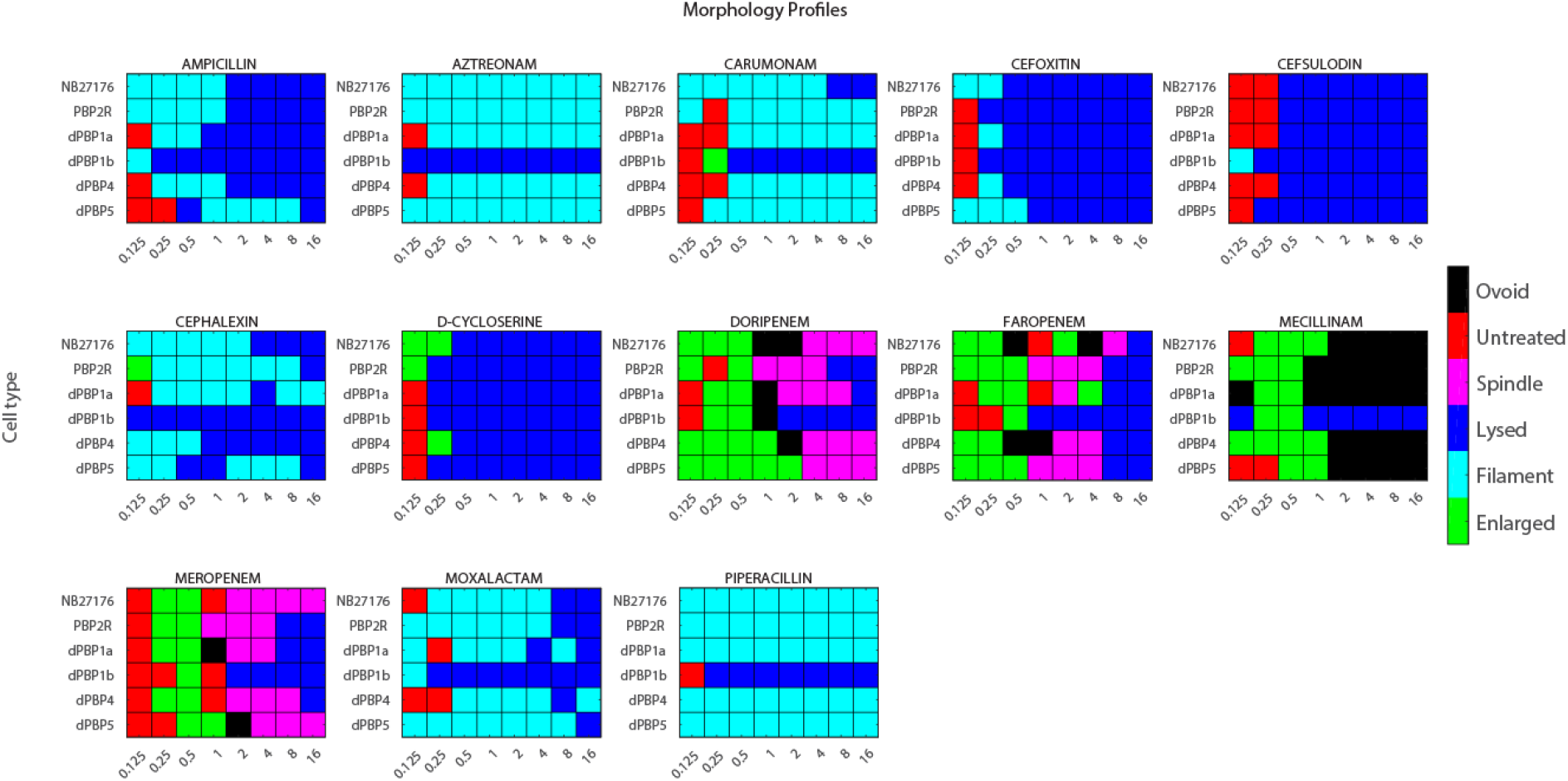
”Tetris plot” visualization of classifications for each dose/strain combination. Morphology scores were averaged over multiple fields of view obtained from at least two independent experiments.

### Morphology profiles are consistent with genetic and microbiological literature

To build confidence in our morphological classifications, we compared our results to literature precedent, where some beta-lactams have been documented with relative specificity for a particular PBP through combined genetic and binding assays. Monobactams such as aztreonam and carumonam reproducibly exhibit strong affinity for PBP3^53,56^ in published gel competition data, and treatment with aztreonam causes a morphology similar to that caused by partial loss of function of PBP3.^16^ Our own gel-binding experiments also reproduced this observation **(Supplementary Figure 1**, **Supplementary Table 2).** Furthermore, cells lacking PBP1b are known to lyse when treated with aztreonam.^43,57^ Consistent with these results, our classifier indicated that treatment with carumonam and aztreonam caused filamentation in all cell lines, with the exception of *E. coli*-Δ*pbp1b*, where it described lysis. Mecillinam has been previously documented as a specific PBP2 inhibitor and is known to cause wild-type cells to adopt an ovoid morphology, identical to that caused by loss of PBP2 activity.^58^ Our classifier made similar predictions, though at low concentrations cells were classified with an enlarged phenotype, which we hypothesize to be a transitory morphology as cells round into the more recognizable ovoid shape. Cells lacking PBP1b were disposed to lysis instead of rounding, which is consistent with past studies of PBP2 and mecillinam.^43,57^ A third molecule, cefsulodin, has been previously demonstrated to bi-specifically inhibit PBP1a and PBP1b, causing lysis in WT cells.^18,52,59–62^ The model recognized similar morphologies in our images. We also measured the effects of D-cycloserine treatment, which inhibits an earlier step in cell wall biosynthesis than beta-lactams. Here, we also observed lysis under all conditions, consistent with previous reports.^63-65^

Having established that our classifier’s morphological predictions were consistent with historical studies, we studied the morphological effects of treatments with more promiscuous classes of beta-lactam antibiotics. The carbapenem and penam classes of beta-lactam have previously been shown to bind to multiple PBPs in binding experiments.^6,8^ Furthermore, molecules such as doripenem and meropenem induce a distinctive phenotype in WT *E. coli* exemplified by becoming more ovular and adopting a spindle morphology.^51,60^ Because this phenotype resembles co-treatment of mecillinam and aztreonam, others have hypothesized that this phenotype is caused by simultaneous inhibition of PBP2 and PBP3 (**Supplementary Figure 2**).^50^ Gel-binding data also suggests that this class of compounds has some affinity for PBP1a/b (**Supplementary Figure 1**). Accordingly, we observe lysis in cells lacking PBP1a. We also observed that in *E. coli-PBP2*^*R*^, carbapenem treatment at low concentrations induced the spindle phenotype but at higher concentrations induced lysis. This behavior was unique to the carbapenems we tested. Faropenem, a member of the related penem class, has also been observed to induce the spindle morphology,^67,68^ and we observed spindle formation at low concentrations. However, with increasing concentration we observed lysis across all strains. This observation is consistent with the increased affinity faropenem displays for PBP1a/b in the gel-binding assay (**Supplementary Figure 1, Supplementary Table 2**). The other agents we tested, corresponding to the penicillin and cephalosporin classes, induced some degree of filamentation, consistent with PBP3 inhibition, but also a tendency to lyse at low concentrations. This suggests that these molecules also inhibit PBP1a/b in cells. Notably, we also observed some unexpected predictions; for instance: filament classifications for low doses of cefsulodin in cells lacking PBP1b. After examining these images, we observed that these predictions may be resulting from ambiguous predictions from the model.

Having classified all the images in the data set, we also compared the cell lines themselves for morphological differences. Interestingly, untreated cells from the *E. coli-PBP2*^*R*^ strain exhibited a higher tendency to be classified as “enlarged,” suggesting that this allele itself has an effect on cell morphology, even in the absence of compound (**Supplementary Figure 3**). Manual examination of the images supported this conclusion; we observed unusually-shaped cells at a higher rate than normal, though they still represent a minority of all objects in these images. Other cell lines, such as *E. coli-ΔPBP5*, appeared no different from WT cells, consistent with previous reports suggesting that its role in morphology is minimal when other Class C enzymes are present (**Supplementary Figure 3**).^21^

### Inferring PBP-binding spectrum of b-lactams via Hidden Markov Models

To facilitate quantitative comparisons between compounds, we sought to devise a dissimilarity function to compare compound morphology profiles. While pairwise comparisons could be carried out by correlating the morphologies observed at each condition, we reasoned that the morphology *dynamics* observed across concentrations would provide additional information that differentiated compound activity. Additionally, for each condition, instead of considering only the morphology with the largest score, we aimed to carry over all morphology scores into the analysis (cf. workflow **Fig 4a**). The morphology scores encode the *uncertainty* of the morphology prediction and may be used to account for borderline predictions. To describe such sequential data with uncertain measurements, we used the hidden Markov model (HMM) formalism^41^, which supports the calculation of dissimilarity measures between sequences^69^.

**Figure 4.**
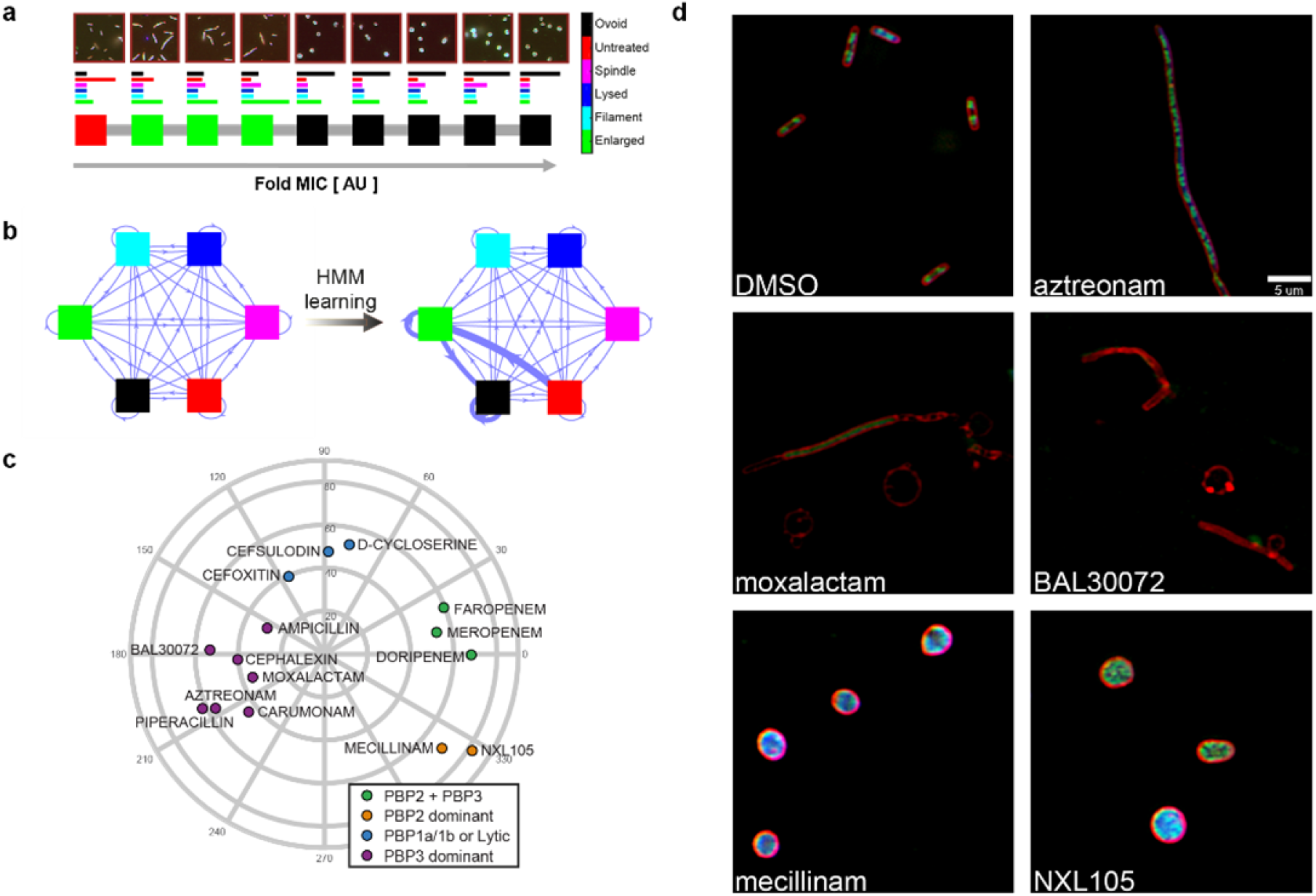
Hidden Markov modelling of bacterial morphology over concentration. (**a**) Images over increasing concentration and corresponding morphology scores, shown as color-coded histograms, derived automatically with a deep neural network. The morphologies with the largest scores at each concentration entail a sequence of morphologies over concentration. (**b**) A graph with nodes representing morphologies and edges representing all possible transitions between morphologies as encapsulated by a hidden Markov model. The width of the edge represents the transition probability between two morphologies. All transition probabilities are derived automatically from the morphology predictions computed by the neural network. The learning process yields transitions with increased probabilitity (wider edges). (**c**) Morphology chart derived from the distances calculated using HMMs that reflects the relationships among different beta-lactam inhibitors. Data points are colored according to the results of a hierarchical clustering algorithm. (**d**) Images corresponding to treatment of WT *E. coli* cells with NXL105 and BAL30072, two recently described PBP inhibitors.

More specifically, for each compound, and for each cell line, we describe the sequence of morphologies observed over increasing MIC folds through an HMM. Within each HMM, we assume that, at a given MIC fold, the bacteria exhibit a certain (hidden) morphology, which is manifested through the corresponding image data and captured through the morphology prediction computed by the neural network. To generate a prediction for a given concentration, the model only takes into consideration the morphology at the immediately prior concentration, that is, we assume that the progression of morphologies over concentration is described by a (first-order) Markov chain. Each HMM is parametrized by the probabilities of transitioning between morphologies, the probabilities of observing a certain morphology while assuming a certain morphology to be the true one, as well as the initial probabilities for each morphology. All parameters are automatically learned from the sequence of morphologies of the corresponding cell line (**Supplementary Methods**). Per compound, we learned the parameters of six HMMs corresponding to the six cell lines in our assay.

Once we determined the parameters of each HMM, we are able to evaluate the probability of observing a sequence of morphologies given the model parameters. To compare two compounds based on their morphological profiles, we interrogate the six HMMs of each compound with the corresponding six sequences of morphologies of the other compound, record the log-probability of observing each sequence, and aggregate these log values into a dissimilarity value (see **Supplementary Methods** for details). Using this strategy, we compare all possible compound pairs, record their dissimilarity values, and store these values into a dissimilarity matrix (**Supplementary Figure 4**). From this dissimilarity matrix, we are able to both group and map compounds onto four clusters in a 2-D space (**Fig. 4c**) that capture the distinct morphological activities of different compounds (**Supplementary Methods**). This *morphology chart* confirms many of the observations made in the beta-lactam literature. In this chart we observe that the reference compounds cefsulodin, aztreonam, and mecillinam, with known distinct PBP preferences, are well separated from one another and are grouped onto different clusters. Cefsulodin is most closely related to D-cycloserine and cefoxitin, which agrees with their propensity to induce lysis. Cefsulodin and cefoxitin both appear to have a high affinity for PBP1a/1b, though cefoxitin appears to have also have affinity for PBP3,^52,70^ which is also reflected by the proximity of this compound to the PBP3 cluster in the chart. Likewise, aztreonam clusters with carumonam, a close structural analogue. Mecillinam, which induces predominantly enlarged and ovoid morphologies across most cell lines, is positioned in an opposite sector in the chart. Similar to mecillinam, the carbapenems also exhibit a prevalent enlarged morphology across most cell lines at lower concentrations. The HMM-based approach however captures the transition between the spindle and lysis morphologies that is unique to these compounds, and thus manages to separate them onto a different neighboring cluster. In this fashion, we see that the dose-dependent morphological dynamics across this panel of strains recapitulates the expected relationships of these molecules based on their historical study using genetic and binding assays. Furthermore, incorporation of all morphology scores into the HMMs suggests relationships that are more accurate than those derived from simple observation of the morphology predictions as depicted in Figure 3; for instance, moxalactam and cephalexin share a similar propensity for filamentation at low doses, followed by lysis at higher concentrations. This is evident from the topology, but not from Figure 3, where lysis is overemphasized in the case of cephalexin.

### Application of the method to recent beta-lactam and diazabicyclooctane analogues

To better understand the prospective value of our method, we profiled two recently described molecules, NXL105 and BAL30072 (**Table 1, Supplementary Table 1, Supplementary Figure 1**). NXL105 is a member of a novel class of beta-lactam mimetic of the diazabicyclooctane class known to inhibit beta-lactamases as well as PBP2 function^71^. BAL30072 is a monocyclic sulfactam bearing a siderophore moiety^72^. These molecules were not part of the training set data. Upon profiling, we observed that NXL105 mapped near mecillinam, consistent with PBP2 inhibition. Accordingly, upon manual observation we observed that the morphology was quite similar to that caused by mecillinam treatment (**Fig. 4c**). This was confirmed internally using the gel-based competition format, and is consistent with the reported behavior of other members of this class (**Supplementary Figure 1**).^73^ Interestingly, upon close examination of the corresponding Tetris plots (**Supplementary Figure 5**), we observed some tendency to classify NXL105-induced morphologies as spindle; however, these appear to be ambiguous predictions that do not substantially affect its placement on the morphology chart. BAL30072, which is structurally related to aztreonam and carumonam, causes morphological changes similar to moxalactam and cephalexin. Our approach accordingly maps BAL30072 in the vicinity of all these compounds. While at low concentrations BAL30072 induces filament formation, higher concentrations resulted in lysis across the cell line panel. This profile, suggestive of PBP1a/b inhibition, is consistent with previous reports on BAL30072^55^ as well as with results obtained from the gel-based binding assay (**Supplementary Figure 1**).

### Systematic comparison to binding data

To more fully understand the correspondence between the HMM-derived beta-lactam relationships and the binding data relationships, we systematically examined both sets of data. In the binding data set, the most potent measured IC50 always suggested the dominant morphology observed by imaging. However, for compounds with broader PBP affinity, the secondary activities measured in the gel-based assay were difficult to interpret. In fact, clustering the compounds based on binding data groups faropenem and meropenem with ampicillin (**Supplementary Figure 6**), and places NXL105 equidistant to aztreonam and mecillinam, which have different mechanisms. In comparison, the morphology-derived relationships are more nuanced, separately grouping specific inhibitors, PBP3+1a/b inhibitors and PBP2+3 inhibitors. Other discrepancies were also observed. For instance, aztreonam and ampicillin exhibited measureable affinity for PBP1A/B and PBP2 respectively. However, imaging offer no evidence of morphologies consistent with these mechanisms.

## Discussion

The PBP family of enzymes is among the oldest of pharmaceutical targets. Despite the wealth of knowledge collected regarding their function and therapeutic relevance, the tools available to study these proteins at high throughput are limited. Here, we describe a high-content imaging approach for studying the cellular effects of PBP inhibition. This approach can be carried out using plate-based, automated microscopes; we profiled compounds in dose-response titrations in six different cell lines. This represents a substantial increase in throughput from what can typically be achieved using gel-based methods. Furthermore, generation of image data provides critical information that supports the elucidation of the mechanism of action of antibiotics.^38,74,75^

To extract morphological information from the data generated by our imaging method, we employ a supervised machine-learning method based on a deep convolutional neural network to rapidly classify full-resolution images into bacterial morphologies. The approach requires no prior object segmentation, which can be difficult for filamentous cells, which frequently overlap, or lysed cells, which lack a uniform shape. To build confidence in our image analysis approach, we compared the identified morphologies with data collected from literature. We confirmed key observations regarding the PBP interaction network, including the synthetic lytic activity of PBP1a and 1b and the tendency for loss of PBP1b to sensitize cells to PBP2- or PBP3-mediated lysis.

One limitation of our supervised machine learning approach is that we have limited our consideration to only six pre-selected phenotypes. Morphologies dissimilar to these six would score poorly in all classes and may be difficult to identify from this analysis. Nonetheless, given the extensive PBP literature, we believe there is justification for prioritizing these morphologies, which are critical to the understanding of PBP function. Another limitation is that compounds inducing similar morphologies through fundamentally different mechanisms—for instance, lysis—would be impossible to separate from PBP inhibitors. We observe this in the case of D-cycloserine and cefsulodin, which both inhibit cell wall biosynthesis, but only cefsulodin is known to inhibit PBPs.

In analyzing the large volume of data resulting from these experiments, we observed that the morphological dynamics across doses and cell lines were meaningful. To capture these data in a single quantitative framework we introduced an approach whereby the sequence of morphological classifications observed across doses could be used as inputs to an HMM—derived entirely from morphological data without any user interaction—with one HMM per compound/cell-line pairing. While others have proposed methods for incorporating dose-response information into morphological analysis^75-77^, our approach, to our knowledge, is the first to utilize the HMM formalism. These HMMs, which capture the context afforded by dose progression, can be quantitatively interrogated to illustrate relationships between different compounds and cell lines. Additionally, they are able to incorporate the prediction strengths of the neural network, which are also key to contextualizing the predictions of the model. Comparisons among compounds, as underpinned by the corresponding HMMs, result in relationships that can be visualized in a 2-dimensional space where the positioning of different beta-lactams correlate with their morphology outcomes. Ultimately, this positioning captures the PBP binding preferences of each compound in all tested doses and cell lines. We compared these predicted preferences to the PBP-binding profile determined using gel-based binding experiment sand found that imaging nearly always agreed with the primary target measured in binding studies, but that discrepancies could be observed when considering secondary target affinities. These secondary binding interactions, such as strong binding affinities for PBP4, have several possible explanations, including: interactions that do not happen in the cell, or manifest outside the 2 hour time period we have captured here, or are ultimately irrelevant to compound mechanism.

The use of HMMs to interpret the dose-dependence of morphological change relies on a predefined morphology universe; consideration of novel morphologies—or a sequence of morphologies—would cause changes in the projected space. Furthermore, this method, like all dose-response experiments, is highly dependent on the choice of concentrations tested. We have chosen to standardize our choice of concentrations relative to the MIC, which provides suitable dynamic range for the conditions tested here, though cells lacking PBP1B are exceedingly sensitive to lysis when treated with PBP2 and PBP3 inhibitors. However, MIC data may not be available for all morphologically active compounds and may be a less useful as a reference point for other mechanisms of antibiotic activity.

Thorough inspection of the morphological changes resulting from treatments with more promiscuous beta-lactams revealed unexpected observations. For example, meropenem and doripenem become prone to lysis in some cell lines. Whereas it should be expected that significant PBP3 inhibition should result in cell lysis in *E. col*i-Δ*pbp1b* cells, it is surprising that lysis was also observed in *E. coli-PBP2*^*R*^. We hypothesize that this morphological outcome could result from a specific combination of PBP inhibition in the cell, perhaps in combination with changes in the free concentration of PBP inhibitors stemming from changes in target concentration or affinities.

To further test the performance of our model, we evaluated how recent PBP inhibitors related to the older beta-lactams. NXL105 has been described as a specific inhibitor of PBP2, and indeed, occupies a space in the projection very near mecillinam. BAL30072, a monobactam which was hypothesized to inhibit PBP3 in addition to PBP1a/b, is close to cephalexin and moxalactam, which exhibit similar morphological patterns. It is intriguing to consider what chemical features of BAL30072 allow it to achieve this activity, since it shares the same scaffold with aztreonam and carumonam. Careful study of the binding mechanisms of these compounds could yield insights into the chemical features that govern PBP1a/b binding.

In conclusion, we believe this work serves as a proof-of-concept that high-content imaging and large-scale morphological profiling based on machine learning provides a useful addition to the suite of tools available to study PBP biology. In order to encourage subsequent work, we have made all images and models available for download. We believe that this type of representation of beta-lactam activities could be of use for evaluating the effects of chemical modification to the beta-lactam scaffold. Continued optimization of the scaffold is likely to continue, and this method could be used to simultaneously optimize features such as beta-lactamase recognition with propensities towards certain mechanisms of action; the observation of both PBP3 and PBP2 target mutations in the clinic suggests that targeting multiple PBPs may be useful. Carbapenems and penicillins appear to be the most promiscuous, but beta-lactamase sensitivity has largely compromised the latter group. The ramifications of other blends of PBP inhibition may have additional *in vivo* implications; perhaps PBP1b inhibition, though not affecting efficacy, may be correlated to molecular pharmacodynamics that could also be explored through a systematic study of the time-dependence of morphological change. Our method allows for these types of features to be easily assessed and incorporated into guiding the optimization of future therapeutics.

## Materials and Methods

### Strains and antibiotics used

*E. coli* strains BW25133, *ΔPBP1a, ΔPBP1b, ΔPBP4*, and *ΔPBP5* were obtained as part of the Keio strain collection^78^ and are available from the Coli Genetic Stock Center (http://cgsc2.biology.yale.edu/). For the *PBP2*^*R*^ strain, the T1718A mutation in mdrA was introduced into E. coli strain NB27079 by recombination^79,45^. See **Supplemental Methods** for more details.

### Gel-based assay

Membrane fractions (1.3mg/mL) were prepared as described elsewhere.^7^ 12 μg of membrane fraction were combined with 1 μL of each compound (1% DMSO final). The reactions were then pre-incubated for 40 min at 37°C, to allow for compound-PBP binding. Following this incubation, 1 µL of 200 μM BOCILLIN was added to each sample and the samples were further incubated at 37°C for 30 min, to allow for BOCILLIN-PBP binding. Each sample was then denatured by the addition of 3 μL 4x NuPAGE LDS Sample Buffer and incubating for 10 min at 70°C. Ten μL of each sample were loaded and separated by SDS-PAGE using 4-12% Tris-Glycine gels (1.0 mm X 12 well), at a constant power of 20 mA for 2 hours. Gels were imaged using a Typhoon 9400 (GE Healthcare Life Sciences) gel imager. Gel bands were quantitated using ImageQuant v5.2 software (Molecular Dynamics). For each band pixel density was manually measured and values were reported as a percentage of bocillin binding.

### Antibacterial activity testing

Antibacterial activity was assessed using a broth microdilution assay following the recommended methodology of the Clinical and Laboratory Standards Institute (CLSI).^80^ In brief, fresh bacterial overnight colony growth was resuspended in sterile saline, adjusted to a 0.5 McFarland turbidity standard and then diluted 1:200 into CAMHB to yield a final target inoculum of 5×10^5^ colony-forming units (CFU)/mL. Two-fold serial dilutions of compounds were prepared in 100% dimethyl sulfoxide (DMSO) at 100-fold the highest final assay concentration; the resulting dilution series of compounds were diluted 1:10 with sterile water. Assay microtiter plates, which contained 10 µL of 10-fold final concentration of compound per well, were inoculated with a volume of 90 µL of bacterial inoculum, sealed in a plastic bag to prevent moisture loss and incubated for 20 hours at 35°C in ambient air. Following incubation, assay plates were monitored for bacterial growth with a SPECTRAmax380 microtiter plate reader (Molecular Devices, Sunnyvale, CA) at 600 nm, as well as by visual observation with a reading mirror. The MIC is defined as the lowest concentration of antibiotic at which the visible growth of the organism is completely inhibited. Performance of the assay was monitored by testing gatifloxacin against laboratory quality control strains in accordance with guidelines of the CLSI^81^

### Imaging assay

The evening before the preparation of sample for imaging, an overnight culture of E.coli strains were made by inoculating bacteria from a frozen glycerol stock into 5 ml of cation-adjusted MHBII medium in a 14ml polypropylene round bottom Falcon tube. The pre-culture was then incubated overnight at 37 °C with orbital shaking at 220 rpm. The next morning, a subculture was prepared by diluting an overnight culture with fresh MHBII medium to an OD600 of 0.05. The culture was then outgrown to OD600 of 0.2. Cells were then diluted to an OD600 of 0.01 and 99ul/well of cells were plated into compound ready assay (96 well glass bottom plate). Cells were incubated at 37°C, 90% humidity for 2 hours. After 2 hours of incubation, cells were washed two times with Hank’s Balanced Salt solution. Cells then were fixed by the addition of formaldehyde and glutaraldehyde to 2.5% and 0.04% (final), respectively. Cells were simultaneously stained with 5ug/ml of FM4-64fx, 5ug/ml of Hoechst, and 0.1uM of Syto9 (final). Fixation was carried out for 30 min at room temperature in the dark, after which the cells were washed two times with Hank’s Balanced Salt solution. Plate was sealed with black vinyl film and imaged using an ImageXpress XLS wide-field imager with a 100X 0.85 NA air objective.

### Morphology prediction with deep neural networks

To automatically classify images displaying multiple bacteria into six morphology categories, we used a multi-scale convolutional neural network (M-CNN) model^40^. The parameters of the M-CNN model were optimized automatically by applying the stochastic gradient descent algorithm to 129 morphology-annotated images (**Supplementary Data**) plus images coming from a data augmentation strategy; further learning details are described elsewhere^40^. Once trained, the M-CNN approach takes as input a full-resolution three-channel fluorescence microscopy image and computes as output a probability score for each of the six bacterial morphologies.

The probability scores of images belonging to the same replicate well are collapsed onto a single set of probability scores by taking the median of the scores of the same morphology across images. Likewise, the resulting probability scores of replicate wells of a given experimental condition are summarized onto a single set of probability scores by taking the median across replicates followed by normalization so that scores added up to unity (or 100%). The morphology with the largest score was taken as the morphology prediction for the corresponding experimental condition.

## Acknowledgments

We would like to thank Jennifer Leeds, Tsuyoshi Uehara, Laura McDowell, Phil Arnold and Peter Skewes-Cox for fruitful discussions.

## Author contributions

BF initiated the study. HC conducted the experiments and collected data. DR carried out gel-based competition experiments. CL ran MIC experiments. RLS synthesized the NXL105 compound. SR made the PBP2R strain. WJG designed the computational methods. IH set up the deep learning framework. WJG, XZ, and BF conducted the analysis. WJG and BF wrote the manuscript. All authors read and approved the final version of the manuscript.

## Supplementary Material

**Supplementary Figure 1**: Bocillin competition data across different classes of PBP inhibitors.

**Supplementary Figure 2**: Combination of PBP3 and PBP2 inhibitors produces a morphology that resembles meropenem treatment

**Supplementary Figure 3**: Comparison of untreated cell morphologies classified using the M-CNN classifier.

**Supplementary Figure 4**: HMM-based dissimilarities among compounds.

**Supplementary Figure 5**: Tetris plot summary of morphological changes caused BAL30072 and NXL105.

**Supplementary Figure 6**: Gel-based clustering.

**Supplementary Table 1**: MICs of B-lactams across cell lines

**Supplementary Table 2:** Percent Bocillin blockage at MIC of each compound.

**Supplementary Methods**: Description of computational methods and strain generation.

**Supplementary Software**: Neural network model as well as HMM code.

**Supplementary Data**: Images and network predictions

